# Development and Characterization of 3D Printable Caprine derived dECM/DAS Crosslinked Hydrogels for Tissue Engineering Applications

**DOI:** 10.1101/2025.06.16.659462

**Authors:** Thirumalai Deepak, Lakshminath Kundanati

**Author notes:** Corresponding author: Dr. Lakshminath Kundanati Mailing address: Assistant Professor, Department of Applied Mechanics and Bio-Medical Engineering, Indian Institute of Technology Madras, Tamilnadu, India. Pin: 600036.

## Abstract

The current hydrogels and crosslinkers used in tissue engineering often suffer from limited bioactivity and insufficient mechanical strength, which hinders effective tissue regeneration. To overcome these limitations, this study aims to develop a tunable hydrogel combining caprine-derived decellularized Extracellular Matrix (dECM) and dialdehyde starch (DAS) for potential use in tissue engineering. The dECM bioink was biohybridized with varying concentration of DAS crosslinker to create dECM/DAS hydrogels and its physicochemical and biological properties were characterized. FTIR analysis confirmed DAS’s successful formation through a strong carbonyl peak at 1732 cm^-1^. The rheological characterization indicates the shear-thinning behavior for dECM/DAS. The storage modulus was higher than that of the loss modulus in all the concentrations of DAS crosslinked hydrogel compared to dECM hydrogel. The gelation kinetics show that dECM and DAS crosslinked hydrogel begin to gel at temperatures above 25°C, which is suitable for developing implantable hydrogel. The hemocompatiblity study shows that the dECM and dECM/DAS hydrogels are non-hemolytic at less than 2%. Overall, this naturally derived dECM/DAS hydrogel demonstrates the potential to be utilized for tissue engineering applications.

## 1.0 Introduction

Hydrogels are a 3D polymeric network composed of polymer chains that have tunable physicochemical properties which make it widely accepted in the field of tissue engineering applications^1^. The decellularized extracellular matrix (dECM) is prepared from the tissue or organ by removing the cells from the extracellular matrix (ECM). The dECM is then solubilized with acetic or hydrochloric acid to make the hydrogel^2^. This dECM hydrogel contains natural components like collagen, elastin, glycoproteins, laminin, and fibronectin which have good biocompatibility and biodegradability^3^. These components support cell adhesion and proliferation at donor sites similar to native tissue ^4^. In the recent years, dECM hydrogels have been extensively investigated because of their versatility, less side effects on the target sites, and minimally invasive delivery. This hydrogel can be injected in pre-gel form, gelation of which is triggered by pH, temperature, light, and chemical reactions^9^. Conversely, their lack of biomechanical strength and elasticity, limits their usage in tissue engineering applications.

The addition of crosslinking agents like formaldehyde, glutaraldehyde, and genipin helps in improving this biomechanical strength and structural properties^1^. However, these agents also exhibit unsatisfactory mechanical properties, low biocompatibility, and high cost^5^. The green synthesis of crosslinking agent can be an alternative to chemical crosslinking agents due to their natural availability, biocompatibility, and biodegradability. Starch, being a natural biomaterial; due to its easy availability, biodegradability, thickening agent, gelling agent and good biocompatibility can be converted into dialdehyde starch (DAS), a green crosslinker by a one-step acid hydrolysis process ^6,7^. The dialdehyde starch (DAS) contains an aldehyde group that can easily form an amine bond with ECM proteins through Schiff base crosslinking^8^.

The study aims to investigate whether an optimal concentration of the DAS crosslinker incombination with dECM can be identified to create an 3D implant/scaffold for tissue engineering applications. The decellularized goat pericardium is solubilized into dECM powder and crosslinked with five different concentrations of DAS. The resulting dECM/DAS hydrogels were evaluated to determine which composition exhibits optimal rheological, physicochemical, and hemocompatibility properties.

## 2.0 Materials and Methods

### 2.1 Sample preparation

Fresh goat pericardium was collected from a local slaughterhouse, and the attached fat was gently removed and washed twice with sterile phosphate-buffered saline (PBS) solution, then the tissue was cut into small pieces and stored in a -20°C freezer until decellularization^11^.

### 2.2 Decellularization and characterization of dECM

The goat pericardium was decellularized and protocol optimization was previously reported^12^. Briefly, the native pericardium was immersed in the round bottom flask containing the combination of 0.5% Sodium dodecyl sulfate (SDS) and 0.5% Triton X-100 solution and stirred gently for 6h. Then, the solution was removed and changed with fresh 1% tri-n-butyl phosphate (TnBP) for another 6h, and then, the tissue was washed with 5 mM ethylenediaminetetraacetic acid (EDTA) and 0.02% sodium azide. Then, to remove all the detergents, the dECM were washed with PBS every 2h once until 6 h and the dECM were sterilized using 2% antibiotic/antimycotic solution. The sterility test was performed on the dECM, and the efficiency of sterilization was previously reported ^13^. Finally, the dECM was lyophilized and stored at -20°C.

#### Histology

To verify the decellularization efficiency, the dECM and native pericardium were stained with Hematoxylin and Eosin (H&E) stain^11^. Initially, the sample was fixed in 10% formalin buffered solutions and embedded in paraffin blocks and the tissue was sectioned 6µm thickness in a microtome. Then the sectioned slides were immersed in xylene solution and a series of ethanol solutions and stained with hematoxylin and eosin (H&E). The stained tissue specimen images were captured in Infinity IOX600 series model light microscope.

### 2.3 Synthesis of DAS and characterization

The DAS was synthesized by following one-step acid hydrolysis and oxidation method ^6^. Briefly, 50mL of 0.6M HCL was added to the round bottom flask and 2.94gm of sodium periodate was stirred for 15min in a magnetic stirrer. 2.0 gm of corn starch was added to the reaction mixture, and it was stirred again for 2hrs at 37C. After 2 hrs of incubation, the obtained white powder was washed with distilled water three times and once washed with acetone to avoid starch coagulation. Then, the obtained DAS was dried in an oven at 50°C for 24hrs.

#### FTIR

Fourier transform infrared (FTIR) was performed to identify the chemical functional group of DAS (Perkin Elmer FTIR spectrometer, Germany). FTIR spectroscopy was recorded in the range of 4000–500 cm^-1^ with an ATR mode. FTIR samples of the synthesized DAS and native starch were prepared by grinding them into a fine powder. The powdered samples were then placed on the ATR crystal and secured using the pressure clamp to ensure good contact. For the dECM and dECM-DAS crosslinked hydrogels, the samples were initially lyophilized and then analyzed in solid form using the same ATR mode.

### 2.4 Preparation of dECM solution

The decellularized extracellular matrix (dECM) powder was solubilized by following the protocol from Pati et al. ^14^. Briefly, the lyophilized dECM was immersed in liquid nitrogen and milled into a fine powder with the help of a cryo mill (Retsch, Germany). The dECM powder weighed a 10:1 ratio of pepsin and was solubilized using 0.5M acetic acid with mild agitation until complete solubilization. After complete solubilization, 1 mL of 10X PBS was added to neutralize the dECM solution and immediately stored at 4°C.

### 2.5 Preparation of dECM/DAS bio-ink

To develop the dECM/DAS, different combinations of bioink was prepared (**Table 1.1).** Initially, the DAS was dissolved in 1M NaOH, and the mixture was sonicated for 1hr at room temperature. Then, the various concentrations of DAS (10, 20, 30, 40, 50%) were added to the solubilized dECM, in a dropwise manner as shown in **Figure 3**. While performing the experiments, the pH of the dECM solution was adjusted to 7.4 by using 10 M NaOH, and the temperature of the reaction was maintained between 4-10°C with the help of an ice pack to avoid the gelation of the sample.

**Table 1.1.** Preparations of dECM/DAS, the different combinations of bio-ink.

| dECM(μl) | DAS(μl) | DAS(%) | Name |
| --- | --- | --- | --- |
| 500 | 0 | 0 | dECM |
| 450 | 50 | 10 | DAS_dECM_10% |
| 400 | 100 | 20 | DAS_dECM_20% |
| 350 | 150 | 30 | DAS_dECM_30% |
| 300 | 200 | 40 | DAS_dECM_40% |
| 250 | 250 | 50 | DAS_dECM_50% |

### 2.6 Rheological characterization of dECM/DAS bio-ink

The rheological characterization was performed on the prepared combinations of dECM/DAS to determine the viscosity, frequency-dependent storage modulus (G’) loss modulus (G”) and gelation kinetics^15,16^. The rheometer (MCR302, Anton Paar, Austria) used in the experiments is equipped with a 25 mm parallel plate with a gap of 0.55mm. To calculate the storage and loss modulus, strain sweeps were performed in the frequency range 0.1-10 Hz. The viscosity of the DAS with DCM mixture was measured at 0.01 to 1000 S^-1^ shear rates to determine the shear-thinning property. The *K* and *n* value of dECM and DAS-dECM were obtained by plotting the graph by shear rate vs. viscosity and the data was fit with Power law equation as mentioned below^15,17^.

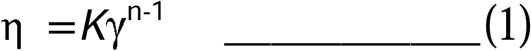

Where, η is the viscosity, γ is the shear rate, *K* is the flow consistency index, *and n* is the power law constant. For the gelation kinetics study, the oscillatory temperature sweep was kept from 4-37°C at a heating rate of 2°C/min.

### 2.7 Swelling study

The swelling study for the dECM hydrogel (n=3) and various concentrations of DAS cross-linked hydrogel (n=3) was performed by immersing the samples in PBS at room temperature for 6 days^18^. For the first day, the wet weight(*W_t_*) of the samples was measured on an hourly basis using a BL220H model weighing balance. After that, every day the wet weight of the sample was measured once for 6 days. Post gelation of hydrogel, the weight of the sample was recorded as the initial weight before immersing in the PBS (*W*_0_). The procedure for the swelling study of the hydrogel is based on the Yeleswarapu et al^19^. The percentage of the swelling (*S*) was calculated by

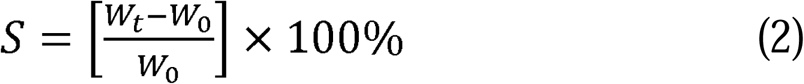

### 2.8 Hemolysis study

The fresh goat’s whole blood was collected from a local slaughter house with 3.2% sodium citrate added as an anticoagulant. The blood was centrifuged at 3000 rpm for 10 minutes and erythrocytes were washed three times with PBS at pH 7.4. 1ml of erythrocytes stock suspension was diluted with 9ml of PBS. Then, the hydrogel sample, washed with PBS was transferred to individual 5ml microfuge tube and incubated with the diluted blood for 1 hr at 37°C. After the incubation time, the samples were centrifuged, and the supernatant was collected. The percentage of hemolysis was calculated by measuring the absorbance of supernatant using microplate reader at a wavelength of 540nm. The PBS was used as a negative control (*A_N_*) (0% hemolysis) and the blood treated with 1% Triton X-100 as a positive control (*A_P_*) (100% hemolysis.) *A_T_* is the absorbance of test sample after incubation with diluted blood. The following formula used to calculate hemolysis

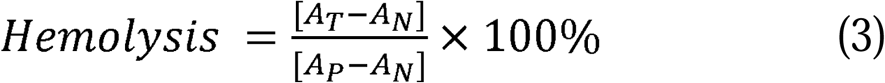

### 2.9 Statistical analysis

All the data were expressed in terms of mean and standard deviation. The one-way ANOVA and Tukey’s multiple comparison tests were performed for the swelling study, hemolysis study and rheological properties using GraphPad Prism (version 9) statistical software.

## 3.0 Results

### 3.1 Characterization

### Histology analysis

Figure:1 shows the cross-section histology of the native and decellularized pericardium stained with H&E staining. When compared to native tissue, the absence of cell nuclei with intact ECM was observed in the decellularized pericardium confirming the complete decellularization.

**Figure 1.**
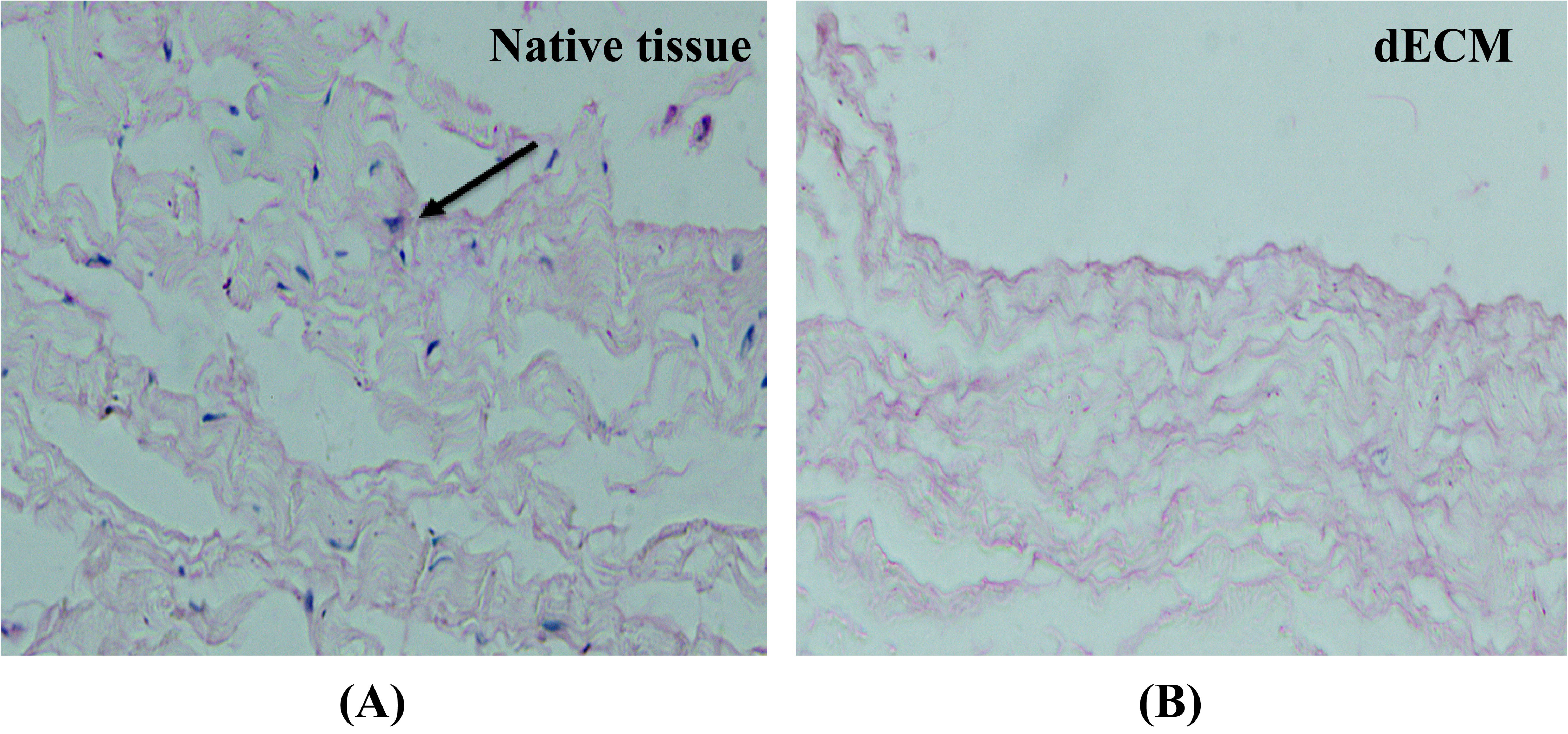
The cross section image of native (A) and decellularized matrix (B). Scale bar 100µm. No signs of cell nuclei or fragments in the dECM compared with native tissue.

Fourier Transform Infrared Spectroscopy (FTIR)The **Figure 2(A)**, represents the comparative FTIR spectrum analysis of the corn starch, and the DAS synthesised from it, while the detailed peak assessments and their corresponding functional groups are listed in **Table 1.2**. The conformational peak for starch was inferred by O-H stretching at 3310 cm^-1^ due to hydrogen bond interaction between the hydroxyl group^6,20^. While, the conformation for DAS was inferred from the peak at 1738 cm^-1^, signifying the presence of aldehyde group. A characteristic peak observed at 2920 and 2931 cm^-1^ in the FTIR spectra for native starch and DAS respectively is attributed to the asymmetrical stretching and vibration of C-H bonds^6^. The characteristic peaks of glycosidic linkages between 1026 and 1000 cm^-1^ respectively indicate the persisting presence of C-O bond in the backbone.

**Table1.2.**
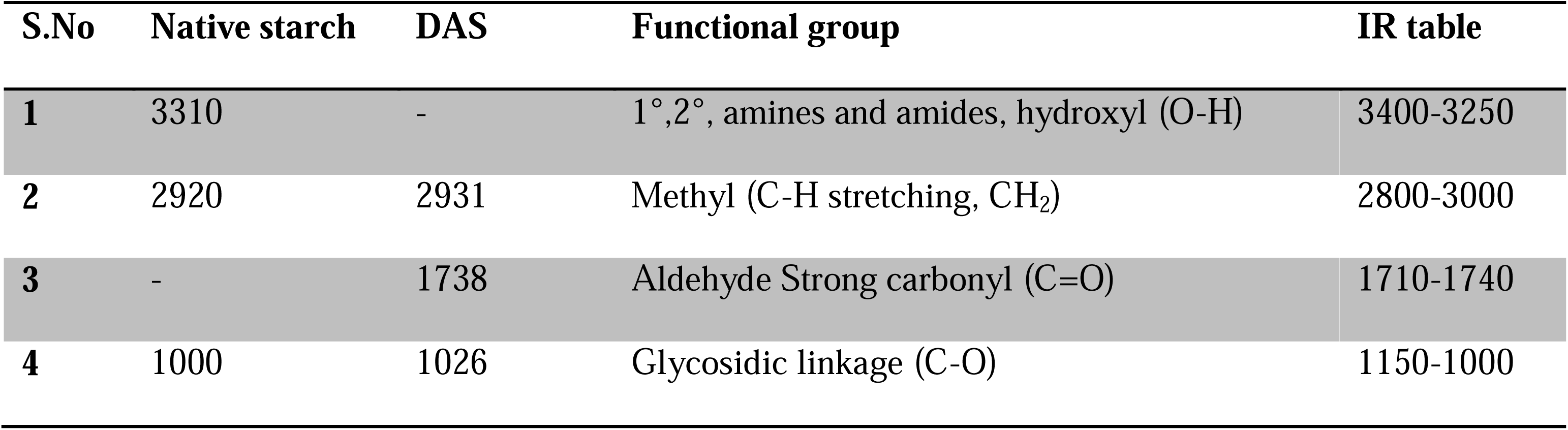
FTIR-ATR spectra peaks of Native starch and DAS crosslinker.

**Figure 2.**
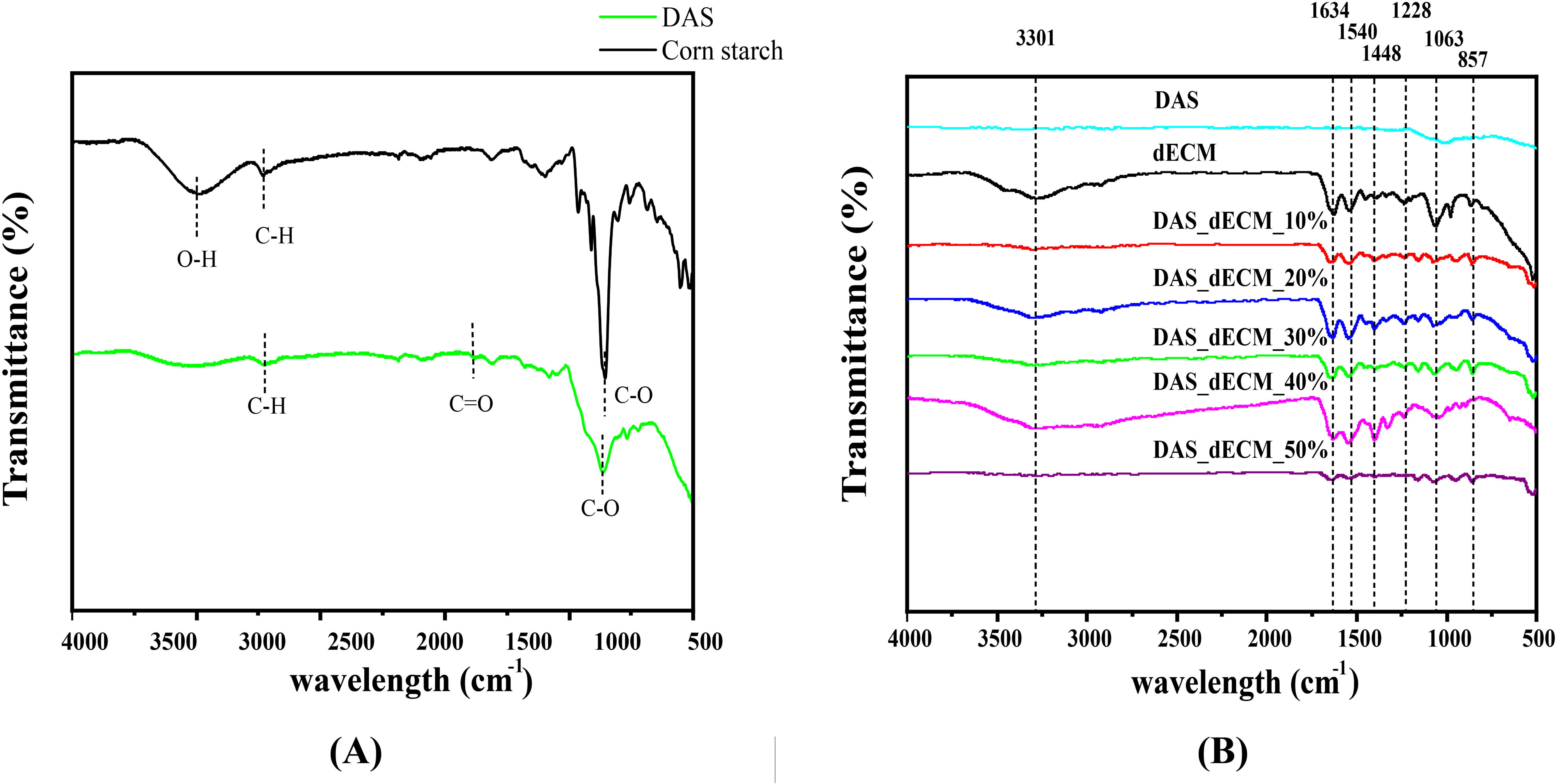
The ATR-FTIR spectroscopy for corn starch and DAS. The peak shows the formation of dialdehyde in corn starch.

**Figure 3.**
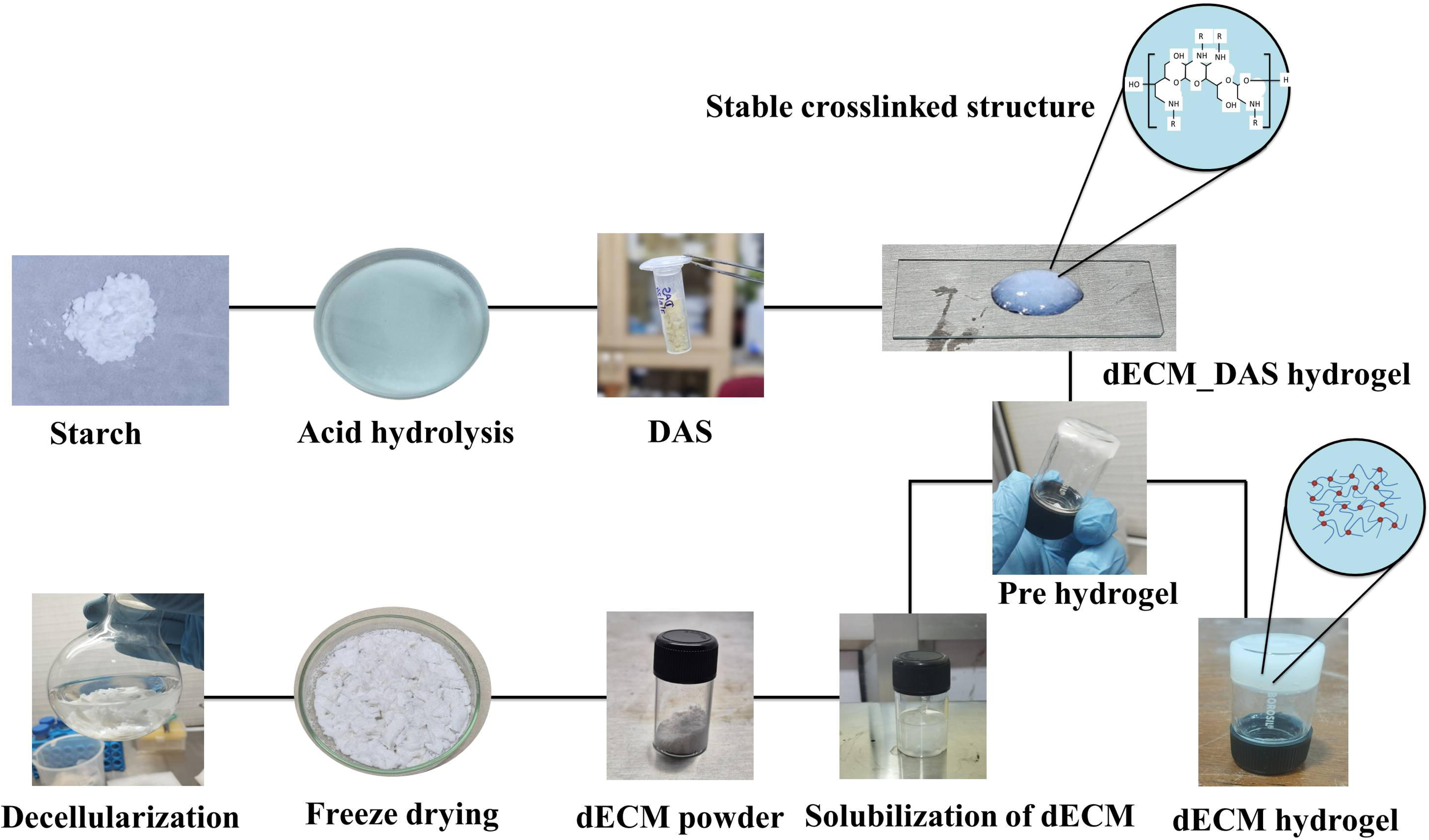
Schematic representation of preparation of dECM and DAS_dECM hydrogel.

dECM hydrogels were produced from caprine pericardium and are composed of collagen, elastin, and glycosaminoglycan (GAG). FTIR analysis showed that O–H stretching, peaks in the range of 3309–3283 cm^-1^ indicating hydrogen bonding within collagen and elastin. The presence of peaks at 1634, 1540, and 1228 cm^-1^ correspond to the amide I, amide II, and amide III bands, respectively, which are attributed to C=O stretching (amide I), N–H bending (amide II), and C–N stretching (amide III) of collagen and elastin^21^. Additionally, the sharp peaks observed between 1450–1400 cm^-1^ are indicative of the presence of GAG content.

In the case of dECM crosslinked DAS, the characteristic peak between 1600–1650 cm^-1^, that is representative of aldehyde (C=O) groups, was not observed. The absence of this peak confirms the consumption of aldehyde groups due to the crosslinking reaction between dECM and DAS. Further, the amide I region overlapped with DAS and slightly shifted from 1634 cm^-1^ to 1624 cm^-1^ which confirms the presence of imine bond structure (C-N) through Schiff-base reaction^7^.

### 3.2 Rheological study

The temperature sweep was performed between 4 to 37°C with an increment of 2°C/min followed by temperature hold at 37°C for 30 minutes (at pH 7.0) as shown in **Figure: 4**. The dECM and DAS crosslinked hydrogel begins to gel at a temperature above 25°C as shown in **Figure: 5(B)**. Through this data, the storage modulus (G’) and the loss modulus (G”) of dECM of the DAS crosslinked hydrogels were determined at 37°C and the results were summarized in **Table: 1.3**. Overall, the G’ value of all the samples were higher than G” signifying the presence of predominant elastic or solid-like behaviour of the sample under the tested conditions.

**Figure 4.**
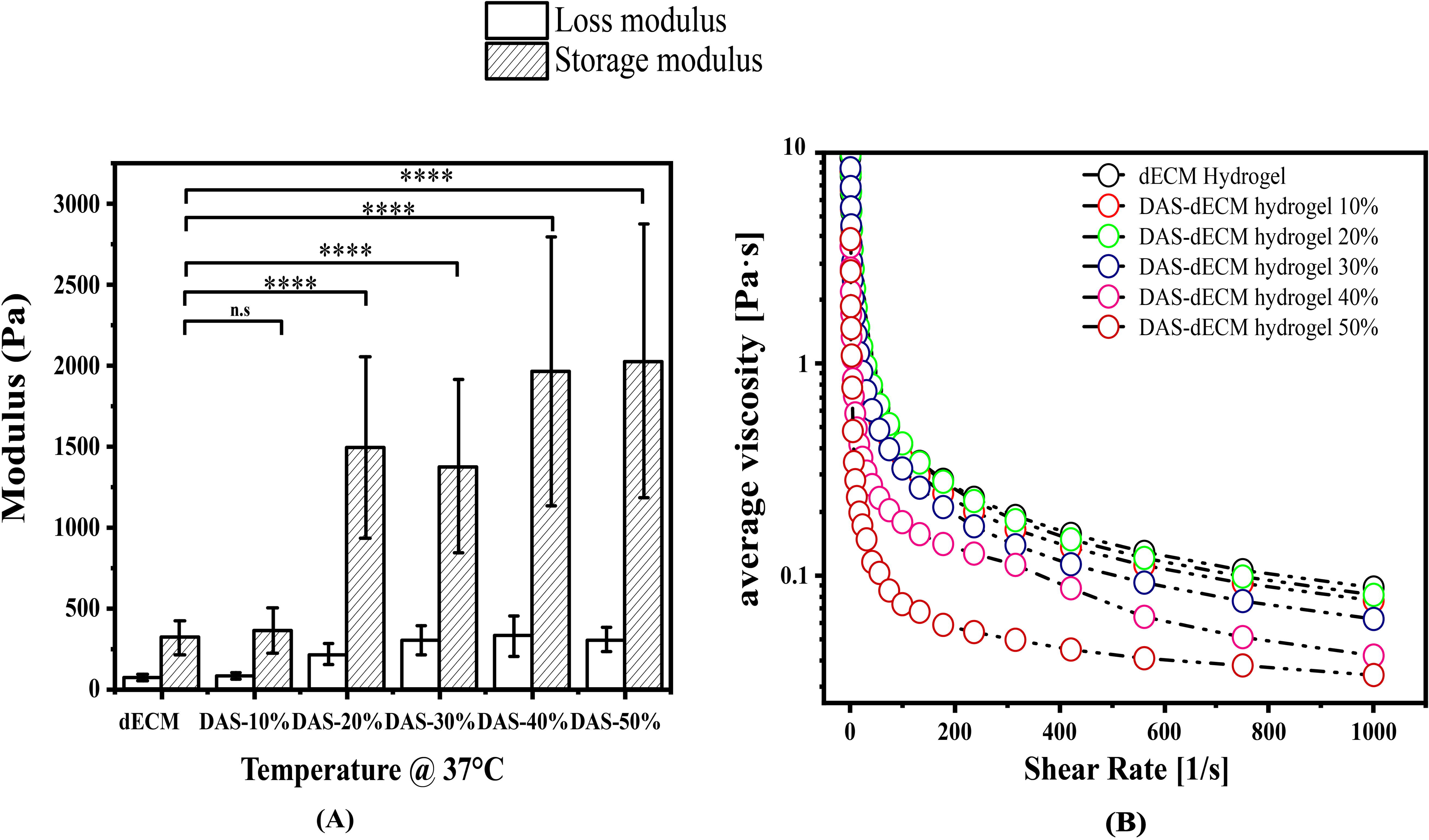
Rheological properties of DAS_dECM and dECM hydrogels. **(A)** Viscosity as a function of shear rate 0.01-1000 s ¹ at 4°C. Measurements were performed using a 25 mm parallel plate geometry with a 0.55 mm gap. **(B)** Strain sweep analysis at 4°C over a frequency range of 0.1-10 Hz. Data are shown as the mean ± standard deviation (n = 3 per group, *p < 0.05, **p < 0.01, ****p < 0.001).

**Figure 5.**
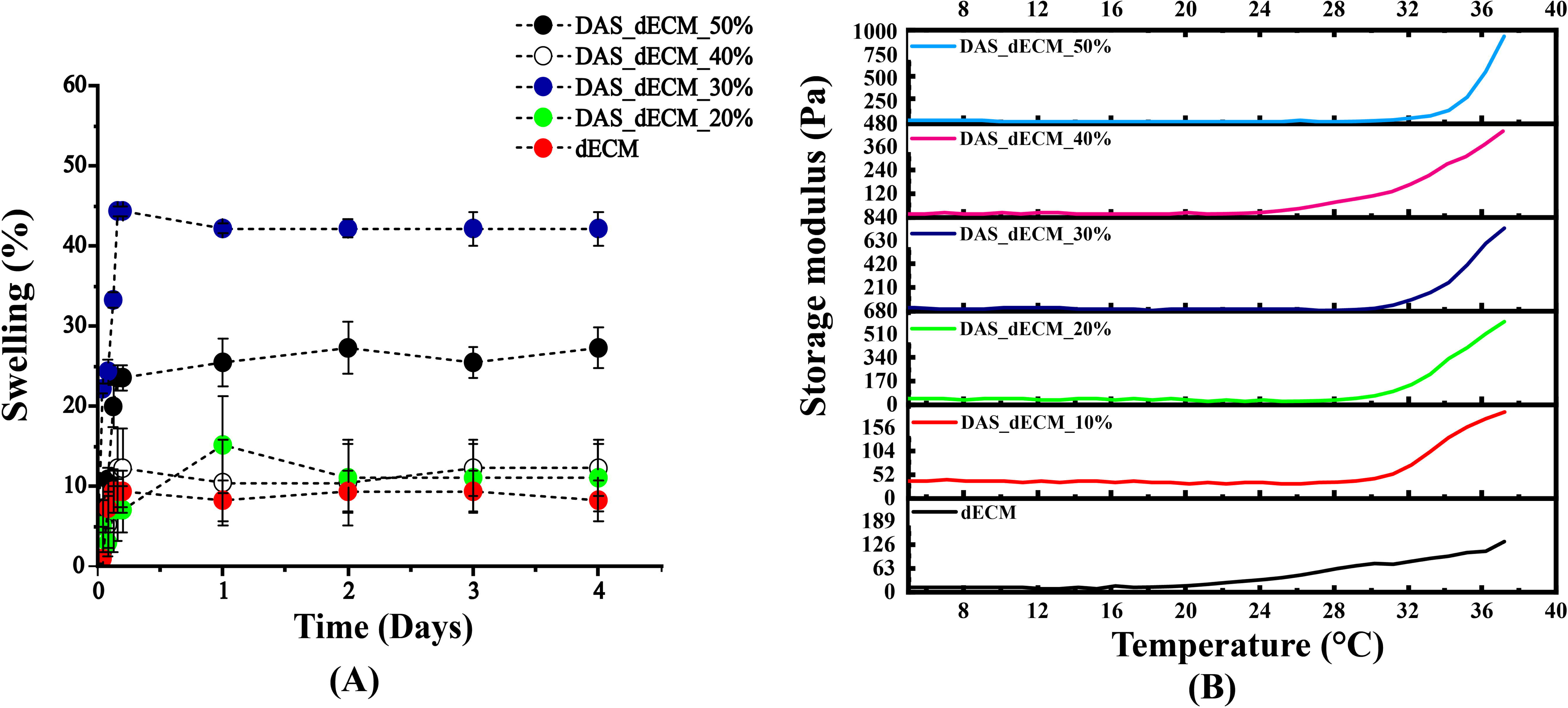
**(A)** Swelling behaviour of various concentrations of DAS_dECM hydrogel and dECM_hydrogel. At the end of 5 Days, all DAS crosslinked hydrogel significantly increased compared to dECM hydrogel. Data represented as mean ± standard deviation; n=3 per group. **(B)** Rheological analysis of dECM and DAS-crosslinked dECM hydrogels during temperature sweep from 4°C to 37°C. All concentrations exhibit a gelation transition above 25°C. Data are shown as the mean ± standard deviation (n = 3 per group, *p < 0.05, **p < 0.01, ****p < 0.001).

**Table1.3.** Rheological analyses of dECM and DAS hydrogel: Viscosity.

| Viscosity | dECM | DAS_ | DAS_ | DAS_ | DAS_ | DAS_ |
| --- | --- | --- | --- | --- | --- | --- |
| @4C |  | dECM_10% | dECM_20% | dECM_30% | dECM_40% | dECM_50% |
| K (Pa) | 13 | 7 | 14 | 11 | 42 | 36 |
| n | 0.42 | 0.19 | 0.30 | 0.36 | 0.11 | 0.02 |

The shear viscosity of the hydrogel was then measured from 0.01 to 1000S^-1^ at 4°C to determine the flow behavior, and the viscosity data was fit with Power law equation, results of which were summarized in **Table:1.4**. The value of n is less than 1, which confirms shear thinning behavior in all dECM hydrogels and no significant (*p*>0.05) difference was observed between them.

**Table 1.4:**
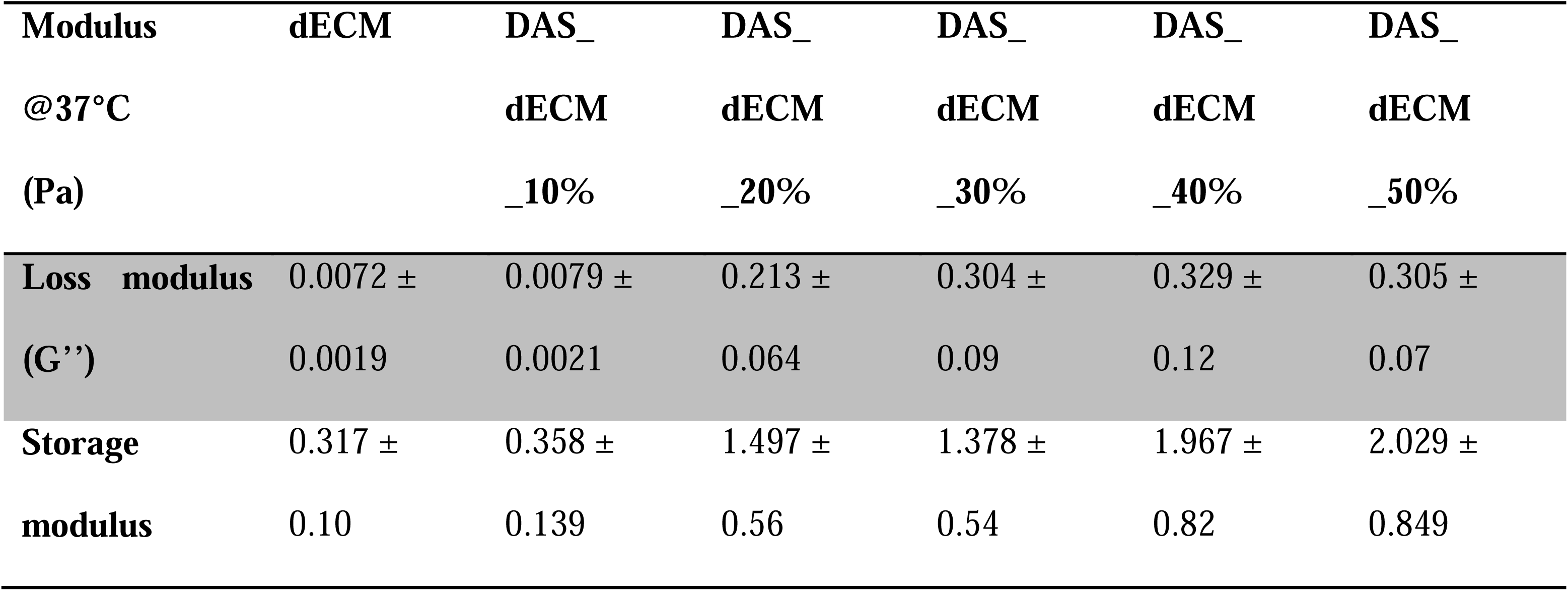
Rheological analysis of dECM and DAS hydrogel: Storage and loss modulus.

| **Modulus** | **dECM** | **DAS_** | **DAS_** | **DAS_** | **DAS_** | **DAS_** |
| --- | --- | --- | --- | --- | --- | --- |
| **@37°C** |  | **dECM** | **dECM** | **dECM** | **dECM** | **dECM** |
| **(Pa)** |  | **_10%** | **_20%** | **_30%** | **_40%** | **_50%** |
| **Loss modulus** | 0.0072 ± | 0.0079 ± | 0.213 ± | 0.304 ± | 0.329 ± | 0.305 ± |
| **(G'')** | 0.0019 | 0.0021 | 0.064 | 0.09 | 0.12 | 0.07 |
| **Storage** | 0.317 ± | 0.358 ± | 1.497 ± | 1.378 ± | 1.967 ± | 2.029 ± |
| **modulus** | 0.10 | 0.139 | 0.56 | 0.54 | 0.82 | 0.849 |

**Table1.5.** Comparison of dECM/DAS hydrogel with existing dECM-based hydrogel for tissue engineering.

| S.No | Material composition and concentration | Method of preparation | Crosslinker | Rheological properties | Swelling ratio | References |
| --- | --- | --- | --- | --- | --- | --- |
| 1 | CdECM +HA | Lyophilization and macro homogenizer | Photocrosslinker-UV 365nm | Storage modulus of the hydrogel increased with increasing UV irradiation | NA | 35 |
| 2 | hUC-dECM | Lyophilization and homogenizer | Thermal gelation at 37°C | A 1.2% dECM concentration exhibits a significantly higher storage modulus $64.04 \pm 6.62$ Pa compared to a 0.6% dECM concentration $28.26 \pm 1.03$ Pa | NA | 36 |
| 3 | HA-NB/DDM | Lyophilization and homogenizer | Photocrosslinker-UV 365nm | G' modulus value of DDM increased with increasing UV irradiation time, indicating that gelation occurred after photo crosslinking. | NA | 37 |
| 4 | dSMM | Lyophilization | Visible light - TEA- Eosin Y- and Thermal gelation at 37°C | Increasing the concentration of eosin from 0.03 mM to 0.5 mM the complex modulus increased by a fold change of 17.6. | Swelling ratio increase 4.2 fold after TEA crosslinking compared to thermal crosslinking | <sup>19</sup> |
| 5 | hdECM | Lyophilization and pulverized by mortar and pestle | Vitamin B2 and Photocrosslinker- 370 nm | 0.02% VB2 crosslinked had the highest complex modulus (10.58 kPa), | NA | <sup>38</sup> |
| 6 | dECM | Lyophilization and cryomill | DAS | Compared to the dECM hydrogel, the DAS crosslinked hydrogel exhibit shear thinning behavior | At the end of 5 Days, all DAS crosslinked groups exhibited significantly swelling than dECM hydrogel. | Present study |

NA- data not available, CdECM-HA-crow scapular cartilage-hyaluronic acid, hUC-dECM- Human Umbilical cord decellularized matrix, HA-NB/DDM-decellularized dermal matrix combined with o-nitrobenzene-modified hyaluronic acid, ASCs-adipose-derived stem cells, TEA- Eosin Y-Triethanolamine, dSMM-decellularized smooth muscle tissue, hdECM-decellularized pig heart tissue, DAS-Dialdehyde starch.

### 3.3 Invitro swelling study

The swelling degree of the dECM and DAS crosslinked hydrogel incubated in PBS is shown in **Figure: 5(A)**. At the end of 5 Days, all crosslinked treatment groups exhibited significant swelling. The swelling percentages were as follows; dECM: 8.3 ± 2.52%; 20% DAS: 11.1 ± 4.16%; 30% DAS: 42.2 ± 2.08%; 40% DAS: 12.3 ± 3.51%; and 50% DAS: 27.3 ± 2.52%. Overall, the swelling behavior of dECM and dECM/DAS hydrogels exhibited a non-linear trend with increasing DAS concentration. The 10% DAS crosslinked hydrogel was unable to retain its shape and was therefore excluded from the graph.

### 3.4 Hemolysis study

According to the biological safety guidelines for biomaterials, a hemolytic index value above 5% is exhibited by hemolytic materials and below 2% by non-hemolytic materials ^22^. The study of the dECM and DAS crosslinked hydrogels exhibited non-hemolytic behavior at a hemolysis value of less than 2% (**Figure 6)** which is well below the acceptable threshold limit. There is no significant difference between the dECM hydrogel and the dECM/DAS crosslinked hydrogel treatment group. This indicates, dECM hydrogel and DAS crosslinking concentrations do not involve any damage to red blood cells confirming good blood compatibility.

**Figure 6.**
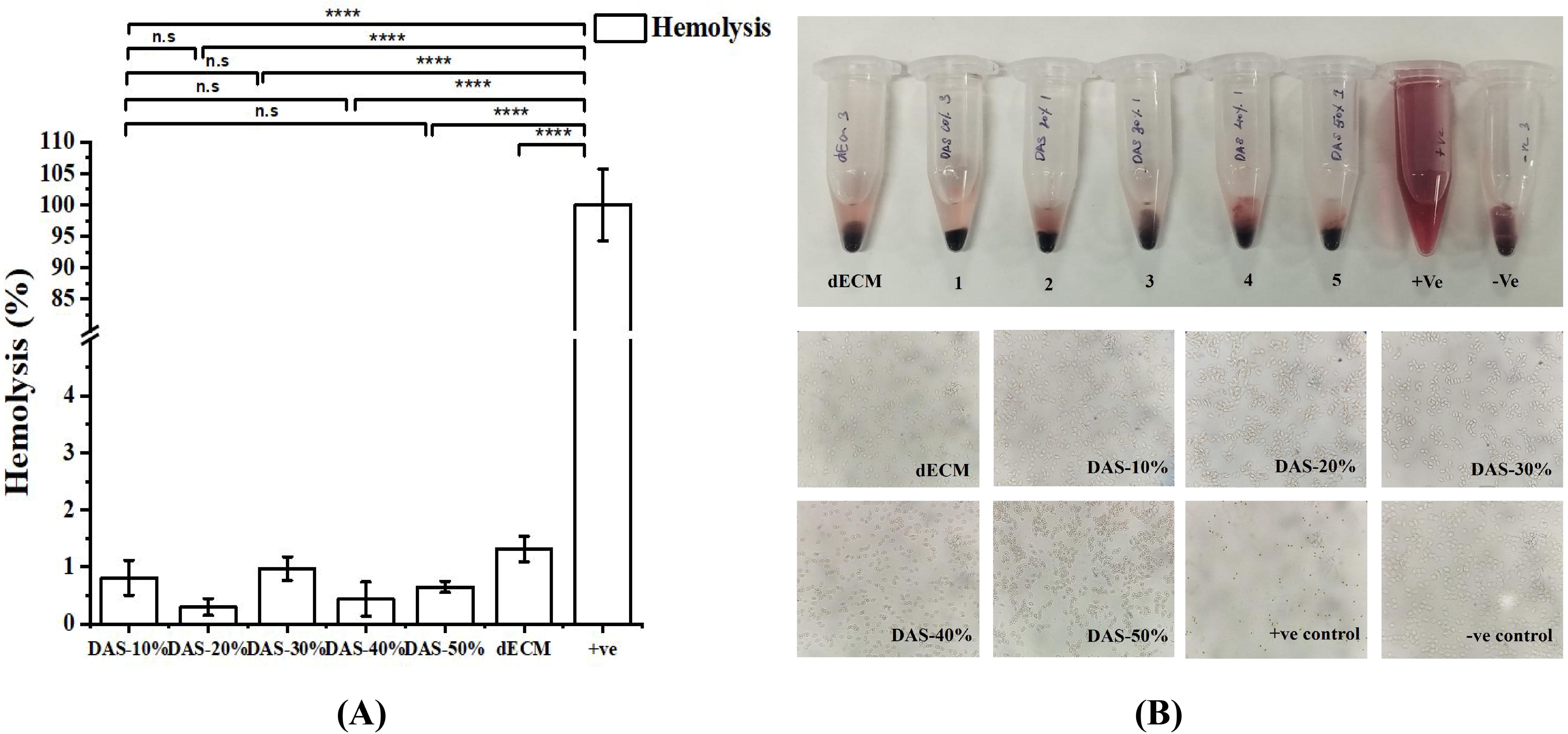
Hemocompatibility analysis of dECM and DAS-crosslinked hydrogels **(A)** Hemolysis obtained for DAS_dECM and dECM hydrogels at various concentrations (10-50%) after incubation with caprine blood. Positive control represents 1% Triton X-100 (complete lysis), negetive control represents PBS (no lysis). **(B)** Microscopic image of blood smear following incubation of DAS_dECM and dECM hydrogels showing intact red blood cells (RBCs) with minimal morphological changes, confirming hemocompatibility. Postive control shows hemolyzed RBCs and negetive control maintains intact RBC morphology. Data are shown as the mean ± standard deviation (n = 5 per group, *p < 0.05, **p < 0.01, ****p < 0.001).

### 4.0 Discussion

The study aimed to create a dECM/DAS bioink for 3D bioprinting by integrating the bioactive properties of solubilized dECM with the natural crosslinking capabilities of DAS, offering a promising approach towards its applications in tissue engineering. Initially, the goat pericardium was decellularized and validated through histology analysis to confirm the cell’s removal and the ECM integrity.

Simultaneously, the natural crosslinker, DAS was synthesized from corn starch through acid hydrolysis. Starch, a natural polymer consisting of amylose and amylopectin linked by α-1,4 and α-1,6 glycosidic bonds, can be converted into DAS through physical, enzymatic, and chemical modification^23^. The FTIR spectroscopy analysis confirmed chemical modification in our study by verifying the formation of aldehyde group in the DAS. Wannous et al. synthesized the DAS using microwave irradiation and performed the FTIR analysis, comparing corn starch with DAS^20^. The results show a new peak at 1702 cm^-1^, corresponding to the stretching vibration of the carbonyl group, which is absent in the starch molecule. This peak indicates the successful synthesis of DAS. Our studyconfirms the formation of DAS by a carbonyl peak at 1738 cm^-1^, consistent with these literature findings and supporting successful DAS synthesis.

Upon incubation of dECM with various DAS concentrations, the pH was adjusted to 7.0 to facilitate Schiff base formation^25^. Crosslinking occurred through both ionic interactions and hydrogen bonding between amine and aldehyde groups, as well as through covalent bonding in Schiff base reaction^7^. FTIR spectra of the crosslinked hydrogels showed that the characteristic amide I band of collagen (∼1624 cm ¹) shifted slightly. This indicates that DAS formed intermolecular interactions with amino acid residues in the collagen triple helix without disrupting collagen secondary structure ^7,20,26^. Thus, the DAS crosslinking approach allows us to obtain stable dECM hydrogel stabilizing the protein-based hydrogels without compromising their native structure. Similarly, Skopinska et al. reported that 10% DAS significantly improved the intermolecular bonding within collagen elastin hydrogels, enhancing mechanical properties, physiochemical properties^26^. Consistent with these findings, our DAS-crosslinked dECM hydrogel also showed improved viscoelastic properties along with rapid gelatinization due to its enhanced physiochemical and biochemical properties ^6,20,24^.

Rheological analysis revealed shear-thinning behavior (n < 1), which is a critical requirement for extrusion-based bioprinting. The printability (K value) in the extrusion-based printer, is in the range of 30-6 ×10^7^ mPas^27^. The K value index increased significantly in DAS-crosslinked samples compared to unmodified dECM, indicating the bioink’s ability to behave as a solid under static conditions and flow under applied stress, which is ideal for extrusion-based printing **(Figure. 4**). In this study, the glass vial inversion method was performed and it showed visual evidence for rapid gelation of the DASdECM and dECM hydrogels, which exhibited a solution-like behavior at low temperature and gel-like behavior at the physiological temperature of 37°C. Real-time monitoring using a rheometer demonstrated that DAS-crosslinked dECM had a higher storage modulus than native dECM bioink at physiological pH. This is due to the covalent bond formation which strengthens the DAS and dECM crosslinking into a three-dimensional network resulting in a more stable hydrogel^10,25,28^.

Hydrogels are inherently hydrophilic and porous, which can compromise mechanical integrity^29,30^. Furthermore, swelling behavior plays a key role in regulating cell adhesion, differentiation, and scaffold performance^29^. Feng et al. classified hydrogels with a swelling ratio of below 150% as non-swelling, which are generally more dimensionally and mechanically stable ^31^. However, hydrogels with more than 150% swelling have high cell migration, and better wound exudate absorption capacity. Hence, an optimal swelling ratio is required based on the intended application of the hydrogel. In this study, DAS crosslinking significantly enhanced the swelling capacity of dECM hydrogels, likely due to increased porosity and the presence of negatively charged proteoglycans, which promote water interaction with polymer chains^31–33^. Notably, the DAS-crosslinked hydrogel maintained a swelling profile within the non-swelling range, making it a promising candidate for applications in tissue engineering and bioelectronics.

When a non-hemocompatible scaffold comes in contact with blood, it results in a series of events, such as hemolysis, protein adsorption, platelet activation, coagulation, and thrombosis. Therefore, hemolytic assay is one of the crucial experiments to confirm hemocompatibility of the scaffold. In this study, dECM and DAS crosslinked hydrogel exhibits non-hemolytic behavior at a hemolysis value of less than 2%. Similar hemolytic results were observed from the decellularized porcine kidney based hydrogel^34^.

## 5.0 Conclusion

This study highlights the potential of using DAS as a natural crosslinker alternative to chemical crosslinker for the development of dECM-based hydrogel. The successful crosslinking of DAS with dECM resulted in a stable, 3D-printable bio-hybrid hydrogel suitable for scaffold fabrication in tissue engineering. FTIR characterization results clearly show that the active bio-hybrid of dECM and DAS, are stable. Rheological parameters such as viscosity, shear stress, gelation kinetics, storage modulus and loss modulus, show that the DAS_dECM hydrogel has shape fidelity, and printability which is suitable for 3D extrusion bioprinting process. Future studies aimed at understanding the biomechanical strength of the DAS_dECM hydrogel and in vitro cytotoxicity, and hemocompatibility testing of the hydrogel.

## Acknowledgement Funding Sources

This project was carried with support from the Department of Applied Mechanics and Biomedical Engineering, NFIG grant of IIT Madras and Ministry of Education, India.

## Statement of ethics

In this study, a cadaveric goat was obtained from the agro-food sector as a donor. As a derivative of food waste, no animal experiments were conducted, and hence, no ethical clearance was required.

## Data availability statement

All the research data presented herein are available on request from the corresponding author.

## Authors contributions

Lakshminath Kundanati (LK) and Thirumalai Deepak (TD) developed the conceptualization, methodology, and research outline. TD conducted the experiment, data acquisition, and manuscript writing—LK supervision, investigation, acquisition of data, validation, writing-reviewing, and editing.

---

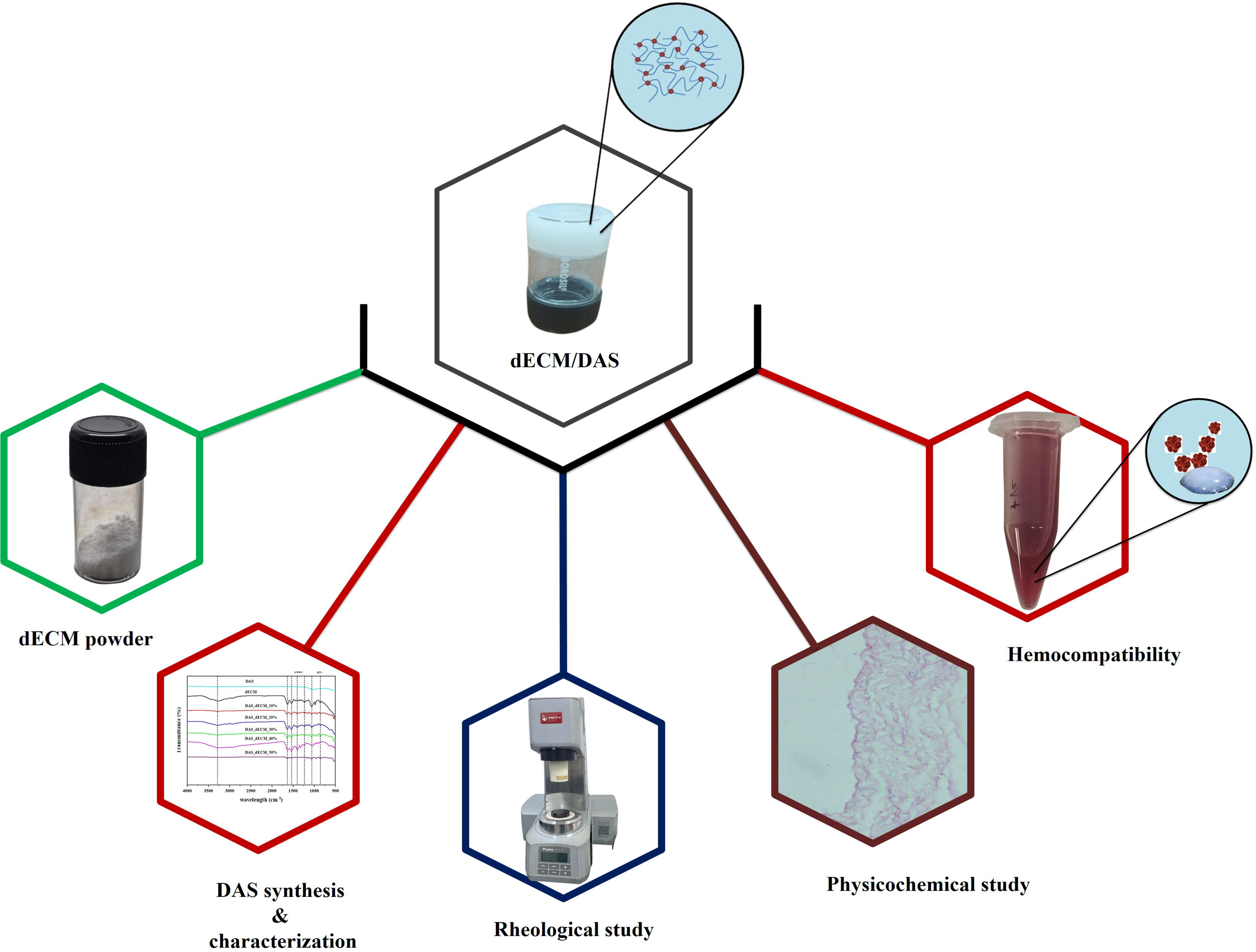

## References

(1) Li, G.; Jiang, Y.; Li, M.; Zhang, W.; Li, Q.; Tang, K. International Journal of Biological Macromolecules Investigation on the Tunable Effect of Oxidized Konjac Glucomannan with Different Molecular Weight on Gelatin-Based Composite Hydrogels. Int. J. Biol. Macromol. 2021, 168, 233–241. 10.1016/j.ijbiomac.2020.12.056.

(2) Zhang, W.; Du, A.; Liu, S.; Lv, M.; Chen, S. Research Progress in Decellularized Extracellular Matrix-Derived Hydrogels. Regen. Ther. 2021, 18, 88–96. 10.1016/j.reth.2021.04.002.

(3) Silva, D. M. da; Barroca, N.; Pinto, S. C.; Semitela, Â.; de Sousa, B. M.; Martins, P. A. D.; Nero, L.; Madarieta, I.; García-Urkia, N.; Fernández-San-Argimiro, F.-J.; Garcia-Lizarribar, A.; Murua, O.; Olalde, B.; Bdikin, I.; Vieira, S. I.; Marques, P. A. A. P. Decellularized Extracellular Matrix-Based 3D Nanofibrous Scaffolds Functionalized with Polydopamine-Reduced Graphene Oxide for Neural Tissue Engineering. Chem. Eng. J. 2023, 472 (July), 144980. 10.1016/j.cej.2023.144980.

(4) Li, H.; Li, P.; Yang, Z.; Gao, C.; Fu, L.; Liao, Z.; Zhao, T.; Cao, F.; Chen, W.; Peng, Y.; Yuan, Z.; Sui, X.; Liu, S.; Guo, Q. Meniscal Regenerative Scaffolds Based on Biopolymers and Polymers: Recent Status and Applications. Front. Cell Dev. Biol. 2021, 9 (July), 1–26. 10.3389/fcell.2021.661802.

(5) Qiao, S.; Peijie, T.; Nan, J. Crosslinking Strategies of Decellularized Extracellular Matrix in Tissue Regeneration. J. Biomed. Mater. Res. - Part A 2024, 112 (5), 640–671. 10.1002/jbm.a.37650.

(6) Zuo, Y.; Liu, W.; Xiao, J.; Zhao, X.; Zhu, Y.; Wu, Y. Preparation and Characterization of Dialdehyde Starch by One-Step Acid Hydrolysis and Oxidation. Int. J. Biol. Macromol. 2017, 103, 1257–1264. 10.1016/j.ijbiomac.2017.05.188.

(7) Cui, T.; Sun, Y.; Wu, Y.; Wang, J.; Ding, Y.; Cheng, J.; Guo, M. Mechanical, Microstructural, and Rheological Characterization of Gelatin-Dialdehyde Starch Hydrogels Constructed by Dual Dynamic Crosslinking. LWT 2022, 161 (March), 113374. 10.1016/j.lwt.2022.113374.

(8) Yang, X.; Liu, G.; Peng, L.; Guo, J.; Tao, L.; Yuan, J.; Chang, C.; Wei, Y.; Zhang, L. Highly Efficient Self Healable and Dual Responsive Cellulose Based Hydrogels for Controlled Release and 3D Cell Culture. Adv. Funct. Mater. 2017, 27 (40), 1–10. 10.1002/adfm.201703174.

(9) Alonso, J. M.; Andrade del Olmo, J.; Perez Gonzalez, R.; Saez-Martinez, V. Injectable Hydrogels: From Laboratory to Industrialization. Polymers (Basel). 2021, 13 (4), 650. 10.3390/polym13040650.

(10) Chae, S.; Lee, S. S.; Choi, Y. J.; Hong, D. H.; Gao, G.; Wang, J. H.; Cho, D. W. 3D Cell-Printing of Biocompatible and Functional Meniscus Constructs Using Meniscus derived Bioink. Biomaterials 2021, 267 (October 2020), 120466. 10.1016/j.biomaterials.2020.120466.

(11) Moffat, D.; Ye, K.; Jin, S. Decellularization for the Retention of Tissue Niches. J. Tissue Eng. 2022, 13. 10.1177/20417314221101151.

(12) Deepak, T.; Bajhaiya, D.; Babu, A. R. Impact of the Different Chemical Based Decellularization Protocols on the Properties of the Caprine Pericardium. Cardiovasc. Eng. Technol. 2024, No. 0123456789. 10.1007/s13239-024-00712-7.

(13) Deepak, T.; Babu, A. R. Investigation of Hemocompatibility and Physicochemical Properties of Acellular Caprine Pericardium for Biomedical Applications. J. Mater. Res. 2023, 38 (7), 1973–1983. 10.1557/s43578-023-00935-9.

(14) Pati, F.; Jang, J.; Ha, D.; Won Kim, S.; Rhie, J.; Shim, J.; Kim, D.; Cho, D. Printing Three-Dimensional Tissue Analogues with Decellularized Extracellular Matrix Bioink. Nat. Commun. 2014, 5 (1), 3935. 10.1038/ncomms4935.

(15) Assad, H.; Assad, A.; Kumar, A. Recent Developments in 3D Bio-Printing and Its Biomedical Applications. Pharmaceutics 2023, 15 (1), 1–45. 10.3390/pharmaceutics15010255.

(16) Habib, M. A.; Khoda, B. Rheological Analysis of Bio-Ink for 3D Bio-Printing Processes. J. Manuf. Process. 2022, 76 (February), 708–718. 10.1016/j.jmapro.2022.02.048.

(17) Basara, G.; Ozcebe, S. G.; Ellis, B. W.; Zorlutuna, P. Tunable Human Myocardium Derived Decellularized Extracellular Matrix for 3D Bioprinting and Cardiac Tissue Engineering. Gels 2021, 7 (2), 70. 10.3390/gels7020070.

(18) Setayeshmehr, M.; Hafeez, S.; van Blitterswijk, C.; Moroni, L.; Mota, C.; Baker, M. B. Bioprinting via a Dual-Gel Bioink Based on Poly(Vinyl Alcohol) and Solubilized Extracellular Matrix towards Cartilage Engineering. Int. J. Mol. Sci. 2021, 22 (8). 10.3390/ijms22083901.

(19) Yeleswarapu, S.; Dash, A.; Chameettachal, S.; Pati, F. 3D Bioprinting of Tissue Constructs Employing Dual Crosslinking of Decellularized Extracellular Matrix Hydrogel. Biomater. Adv. 2023, 152 (May), 213494. 10.1016/j.bioadv.2023.213494.

(20) Wannous, A.; Milaneh, S.; Said, M.; Atassi, Y. New Approach for Starch Dialdehyde Preparation Using Microwave Irradiation for Removal of Heavy Metal Ions from Water. SN Appl. Sci. 2022, 4 (5), 133. 10.1007/s42452-022-05024-w.

(21) Huang, X.; Ding, Y.; Pan, W.; Lu, L.; Jin, R.; Liang, X.; Chang, M.; Wang, Y.; Luo, X. A Comparative Study on Two Types of Porcine Acellular Dermal Matrix Sponges Prepared by Thermal Crosslinking and Thermal-Glutaraldehyde Crosslinking Matrix Microparticles. Front. Bioeng. Biotechnol. 2022, 10 (August), 1–16. 10.3389/fbioe.2022.938798.

(22) Weber, M.; Steinle, H.; Golombek, S.; Hann, L.; Schlensak, C.; Wendel, H. P.; Avci-Adali, M. Blood-Contacting Biomaterials: In Vitro Evaluation of the Hemocompatibility. Front. Bioeng. Biotechnol. 2018, 6, 12. 10.3389/fbioe.2018.00099.

(23) Dong, Y.; Li, Z.; Kong, H.; Ban, X.; Gu, Z.; Zhang, H.; Hong, Y.; Cheng, L.; Li, C. Correlation Analysis of Starch Molecular Structure and Film Properties via Rearrangements of Glycosidic Linkages by 1,4-α-Glucan Branching Enzyme. Carbohydr. Polym. 2025, 348 (PB), 122908. 10.1016/j.carbpol.2024.122908.

(24) Ziegler-Borowska, M.; Wegrzynowska-Drzymalska, K.; Chelminiak-Dudkiewicz, D.; Kowalonek, J.; Kaczmarek, H. Photochemical Reactions in Dialdehyde Starch. Molecules 2018, 23 (12). 10.3390/molecules23123358.

(25) Wang, B.; Barceló, X.; Von Euw, S.; Kelly, D. J. 3D Printing of Mechanically Functional Meniscal Tissue Equivalents Using High Concentration Extracellular Matrix Inks. Mater. Today Bio 2023, 20 (April). 10.1016/j.mtbio.2023.100624.

(26) Skopinska-Wisniewska, J.; Wegrzynowska-Drzymalska, K.; Bajek, A.; Maj, M.; Sionkowska, A. Is Dialdehyde Starch a Valuable Cross-Linking Agent for Collagen/Elastin Based Materials? J. Mater. Sci. Mater. Med. 2016, 27 (4), 67. 10.1007/s10856-016-5677-6.

(27) Bankhede, H. K.; Ganguly, A. Pharmaceutical Polymer-Based Hydrogel Formulations as Prospective Bioinks for 3D Bioprinting Applications: A Step towards Clean Bioprinting. Ann. 3D Print. Med. 2022, 6, 100056. 10.1016/j.stlm.2022.100056.

(28) Elomaa, L.; Almalla, A.; Keshi, E.; Hillebrandt, K. H.; Sauer, I. M.; Weinhart, M. Rise of Tissue- and Species-Specific 3D Bioprinting Based on Decellularized Extracellular Matrix-Derived Bioinks and Bioresins. Biomater. Biosyst. 2023, 12 (June), 100084. 10.1016/j.bbiosy.2023.100084.

(29) Kesharwani, P.; Alexander, A.; Shukla, R.; Jain, S.; Bisht, A.; Kumari, K.; Verma, K.; Sharma, S. Tissue Regeneration Properties of Hydrogels Derived from Biological Macromolecules: A Review. Int. J. Biol. Macromol. 2024, 271 (P2), 132280. 10.1016/j.ijbiomac.2024.132280.

(30) Li, X.; Gong, J. P. Design Principles for Strong and Tough Hydrogels. Nat. Rev. Mater. 2024, 9 (6), 380–398. 10.1038/s41578-024-00672-3.

(31) Feng, W.; Wang, Z. Tailoring the Swelling-Shrinkable Behavior of Hydrogels for Biomedical Applications. Adv. Sci. 2023, 10 (28), 1–41. 10.1002/advs.202303326.

(32) Ghodbane, S. A.; Dunn, M. G. Physical and Mechanical Properties of Cross-Linked Type I Collagen Scaffolds Derived from Bovine, Porcine, and Ovine Tendons. J. Biomed. Mater. Res. - Part A 2016, 104 (11), 2685–2692. 10.1002/jbm.a.35813.

(33) Guo, X.; Park, H.; Temenoff, J. S.; Tabata, Y.; Caplan, A. I.; Mikos, A. G. Effect of Swelling Ratio of Injectable Hydrogel Composites on Chondrogenic Differentiation of Encapsulated Rabbit Marrow Mesenchymal Stem Cells in Vitro. AIChE Annu. Meet. Conf. Proc. 2008, 541–546.

(34) Quinteira, R.; Gimondi, S.; Monteiro, N. O.; Sobreiro-Almeida, R.; Lasagni, L.; Romagnani, P.; Neves, N. M. Decellularized Kidney Extracellular Matrix-Based Hydrogels for Renal Tissue Engineering. Acta Biomater. 2024, 180, 295–307. 10.1016/j.actbio.2024.04.026.

(35) Xu, Y.; Jia, L.; Wang, Z.; Jiang, G.; Zhou, G.; Chen, W.; Chen, R. Injectable Photo-Crosslinking Cartilage Decellularized Extracellular Matrix for Cartilage Tissue Regeneration. Mater. Lett. 2020, 268, 127609. 10.1016/j.matlet.2020.127609.

(36) Xia, W.; Jin, M.; Feng, Z.; Zhang, J.; Rong, Y.; Zhang, Y.; Zhang, S.; Yu, Y.; Yang, H.; Wang, T. Injectable Decellularzied Extracellular Matrix Hydrogel Derived from Human Umbilical Cord: A Novel Perspective to Deal with Refractory Wound via Medical Wastes. Mater. Des. 2023, 229, 111877. 10.1016/j.matdes.2023.111877.

(37) Bo, Q.; Yan, L.; Li, H.; Jia, Z.; Zhan, A.; Chen, J.; Yuan, Z.; Zhang, W.; Gao, B.; Chen, R. Decellularized Dermal Matrix-Based Photo-Crosslinking Hydrogels as a Platform for Delivery of Adipose Derived Stem Cells to Accelerate Cutaneous Wound Healing. Mater. Des. 2020, 196, 109152. 10.1016/j.matdes.2020.109152.

(38) Jang, J.; Kim, T. G.; Kim, B. S.; Kim, S. W.; Kwon, S. M.; Cho, D. W. Tailoring Mechanical Properties of Decellularized Extracellular Matrix Bioink by Vitamin B2-Induced Photo-Crosslinking. Acta Biomater. 2016, 33, 88–95. 10.1016/j.actbio.2016.01.013.

